# BRAin INteractive Sequencing Analysis Tool (BRAIN-SAT); facilitating interactive transcriptome analyses (http://brainsat.eu/)

**DOI:** 10.1101/750000

**Authors:** M.L. Dubbelaar, M.L. Brummer, M. Meijer, B.J.L. Eggen, H.W.G.M. Boddeke

## Abstract

Over the last decade, a large number of glia transcriptome studies has been published. New technologies and platforms have been developed to allow access and interrogation of the published data. The increase in large transcriptomic data sets allows for innovative *in silico* analyses to address biological questions. Here we present BRAIN-SAT, the follow-up of our previous database GOAD, with several new features available on an interactive platform that enables access to recent, high quality bulk and single cell RNA-Seq data. The combination of several functions including gene searches, differential and quantitative expression analysis and a single cell expression analysis feature enables the exploration of published data sets at different levels. These different functionalities can be used for researchers and research companies in the neuroscience field to evaluate and visualize gene expression levels in a set of relevant publications. Here, we present a new platform with easy access to published gene expression studies for data exploration and gene of interest searches.

## 1 Introduction

Due to large number of transcriptomic studies over the last few years and the generation of single cell RNA sequencing data, a vast amount of transcriptome data has become available in various repositories. However, the available datasets are processed with a variety of sequencing techniques and materials, which results in different technological batches and is therefore difficult to compare and combine. Another difficulty is the often unprecise or even incomplete dataset descriptions that make it challenging to compare transcriptomic datasets and to perform meta-analyses. Using the guidelines for data management and storage as outlined in the FAIR concept^1,2^ might lead to improved open accessibility of datasets, enabling re-usage of data.

For the purpose of re-use of data, we previously created the glia open access database (GOAD)^3^ to provide and harmonize previously published high quality glia transcriptome datasets. GOAD contains a collection of selected studies where researchers can evaluate the differences in gene expression by selecting predefined comparisons. To improve GOAD and implement novel functionalities, we developed the brain interactive sequencing analysis tool (BRAIN-SAT) introducing a user-friendly, interactive application to re-analyze published data. This application analyzes raw transcriptome data (bulk and single cell) from available dataset (studies are represented in table 1). For proper harmonization, all datasets in BRAIN-SAT were preprocessed and stored in the same format. These harmonized data sets can then be used for different visualizations to interpret the data (figure 1).

**Figure 1.**
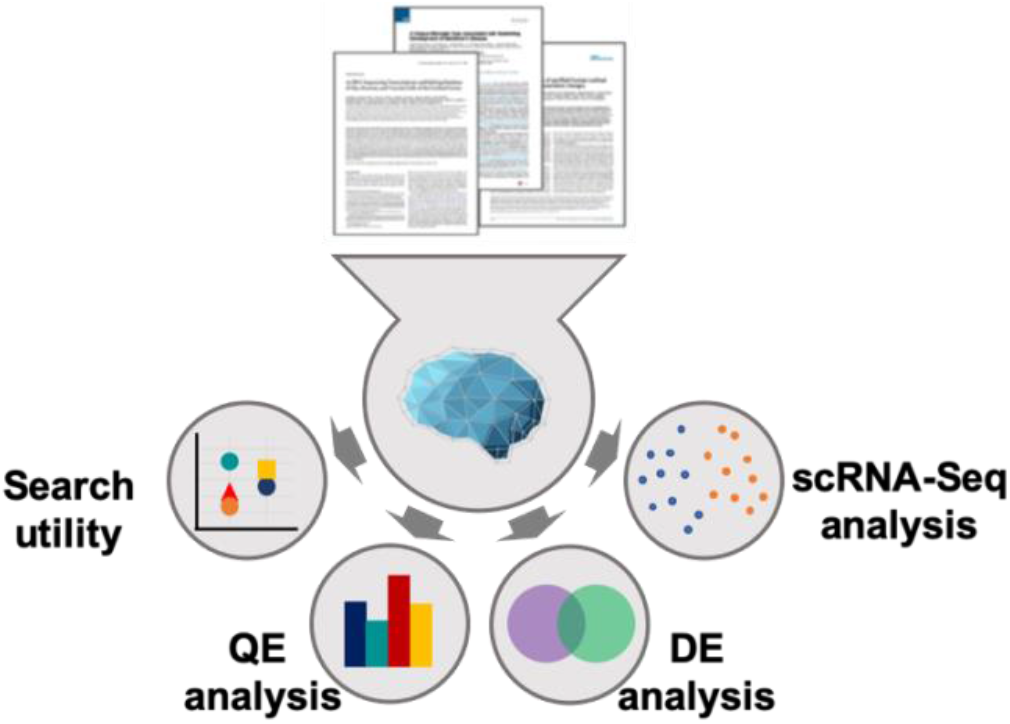
Overview BRAIN-SAT. Raw data of published transcriptome data sets is used as input for BRAIN-SAT where the alignment and quantification are done to obtain the gene counts of the various studies. BRAIN-SAT uses this information of different studies, to process this data with different analysis pipelines to visualize the outcome. The search utility enables the visualization of gene expression levels in different cell types among all available studies. Quantitative expression analysis enables the quantification of gene-expression levels. Differential expression analysis can be performed to determine changes in gene expression between two conditions. The scRNA-Seq analysis offers visualizations in the form of a tSNE plot to depict gene expression differences in the scRNA-Seq studies.

**Table 1.**
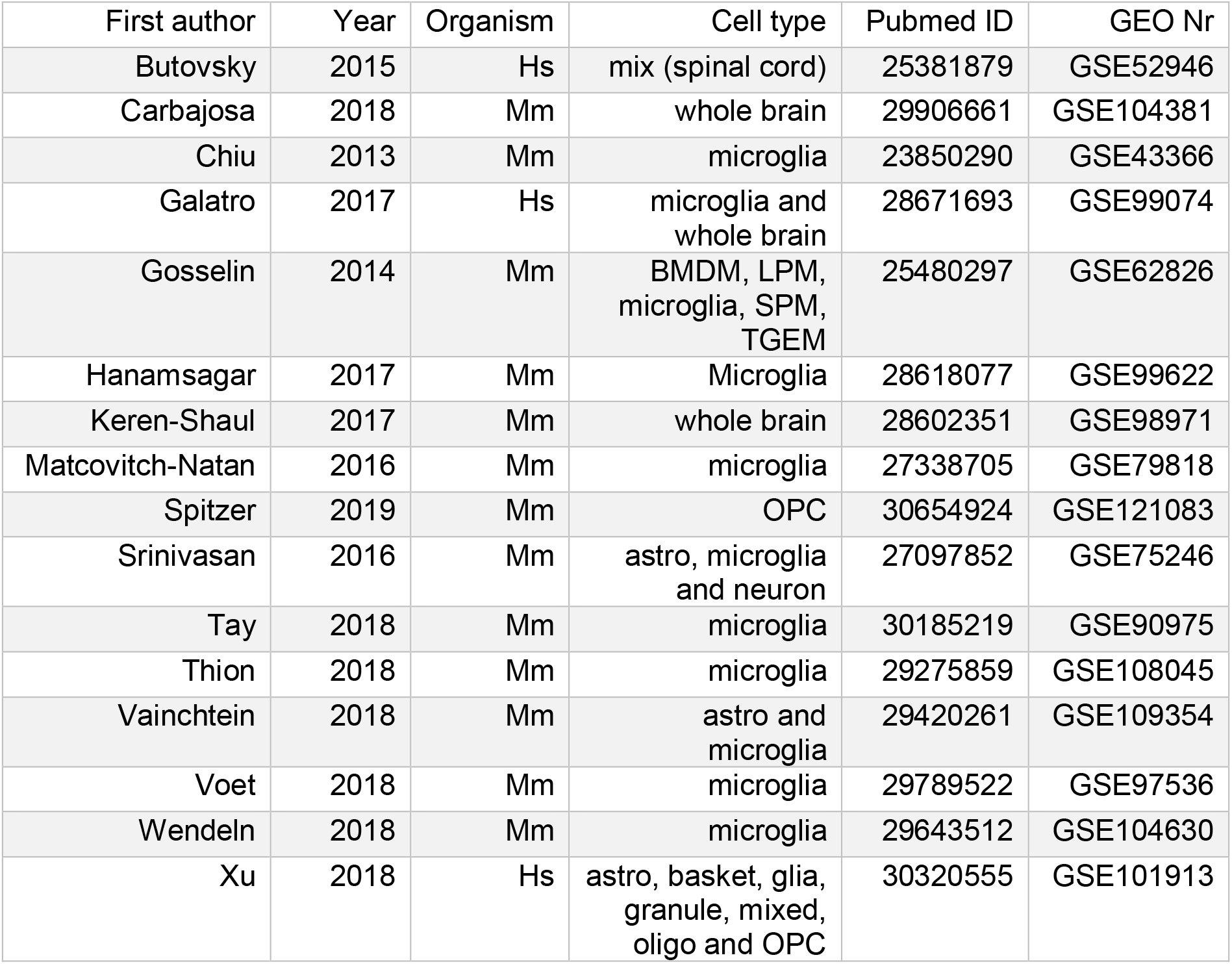
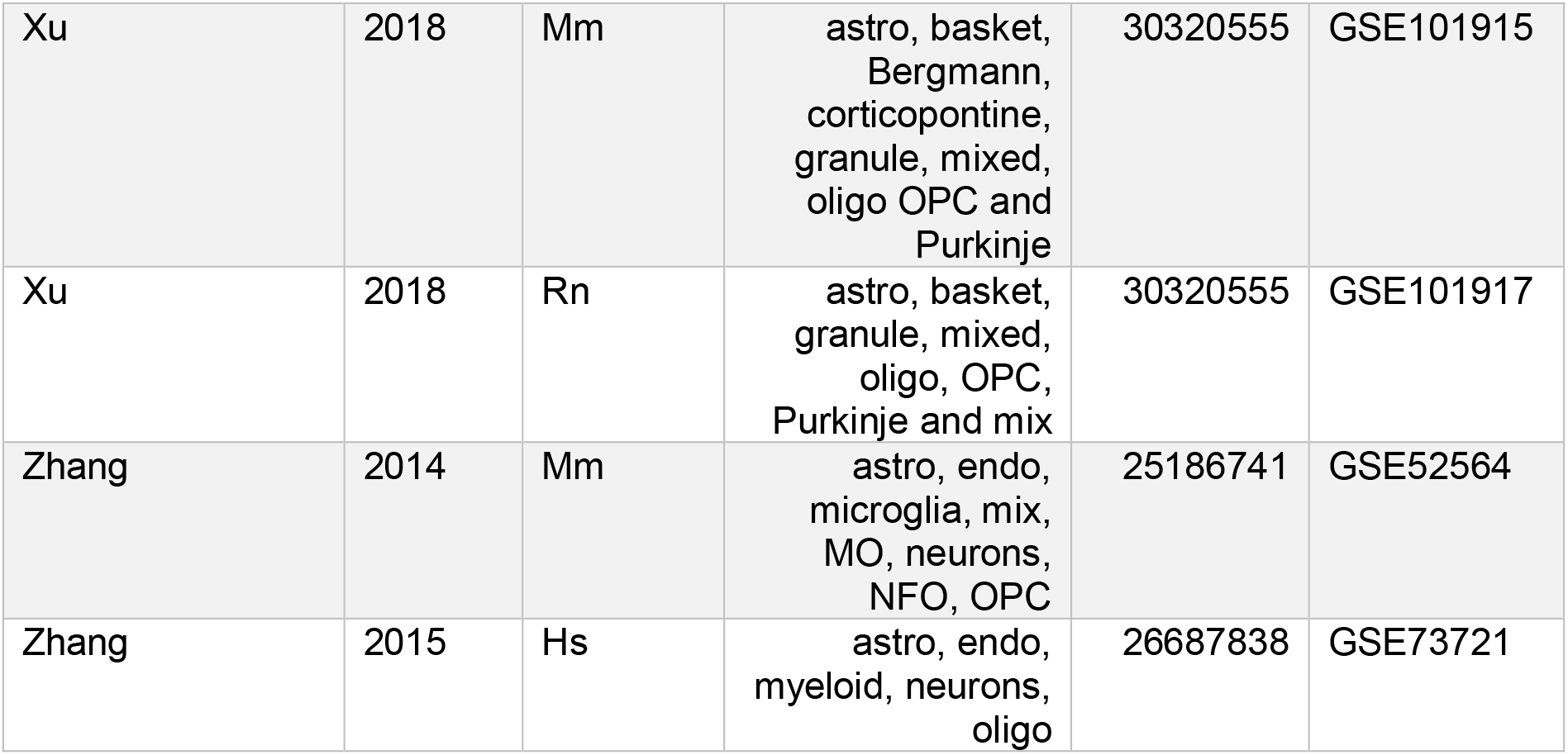
Available studies in BRAIN-SAT

BRAIN-SAT as a successor of GOAD contains more interactive functionalities for transcriptome analysis. The application is built on the database structure of MOLGENIS^4^, where it uses the integrated database and the R application programming interface (API) to perform differential expression analysis and processing of single cell data.

The differential and quantitative expression analysis functions are, like in GOAD, still available but modified into a more user-friendly format. The fast-interactive function is particularly noticeable in the differential expression analysis (DEA) utility where the comparisons rely on raw counts, which are stored in the database. This setup enables us to add new studies faster, since now they only need to be aligned and quantified before the studies are stored in the database. A second interactive part of BRAIN-SAT are the images, which facilitate the outcome of the different analyses, additional information regarding the data can be observed when hovering over the visualization.

An improved feature of BRAIN-SAT is the gene search function. This functionality consists of data values of several studies (cross-study) and organisms in one visualization analysis. The log2(counts per million) of the average expression values per condition were used as data values for this purpose.

In addition, BRAIN-SAT has a new feature i.e. a single cell sequencing analysis function. Several single cell glia studies has been selected to visualize single cell expression data in interactive t-distributed stochastic neighbor embedding (tSNE)^5^ plots where the quantitative expression values of genes are displayed.

## 2 Materials and Methods

### MOLGENIS

MOLecular GENetics Information Systems (MOLGENIS)^4^ is a toolkit that consists of several bioinformatics structures and user interfaces that can be used for managing and processing scientific data. For BRAIN-SAT, several MOLGENIS components were used: the font end, data tables and available scripting tools. The front end represents the BRAIN-SAT layout and is the starting point of various analyses. The MOLGENIS data tables store aligned reads that are used throughout the rest of BRAIN-SAT and can be accessed with the use of the representational state transfer (REST) API. The R API facilitates the interactive analyses for transcriptome data and the JavaScript module enables features that are specific for BRAIN-SAT.

### Preprocessing pipeline

Raw fastq files of bulk RNA-Seq are processed through a standardized pipeline, where fastq files are obtained through the gene expression omnibus (GEO)^6^ or the European Nucleotide Archive (ENA)^7^. Low quality base pairs in the sequence are trimmed. Alignment is performed with the use of HiSat2^8^. Sequences are aligned with the following genomes: human (GRC38), mice and rat (GRCm38). Samtools^9^ and Picard^10^ are used after the alignment, different functions and additional parameters used for the preprocessing steps are explained in table 3.

**Table 3.**
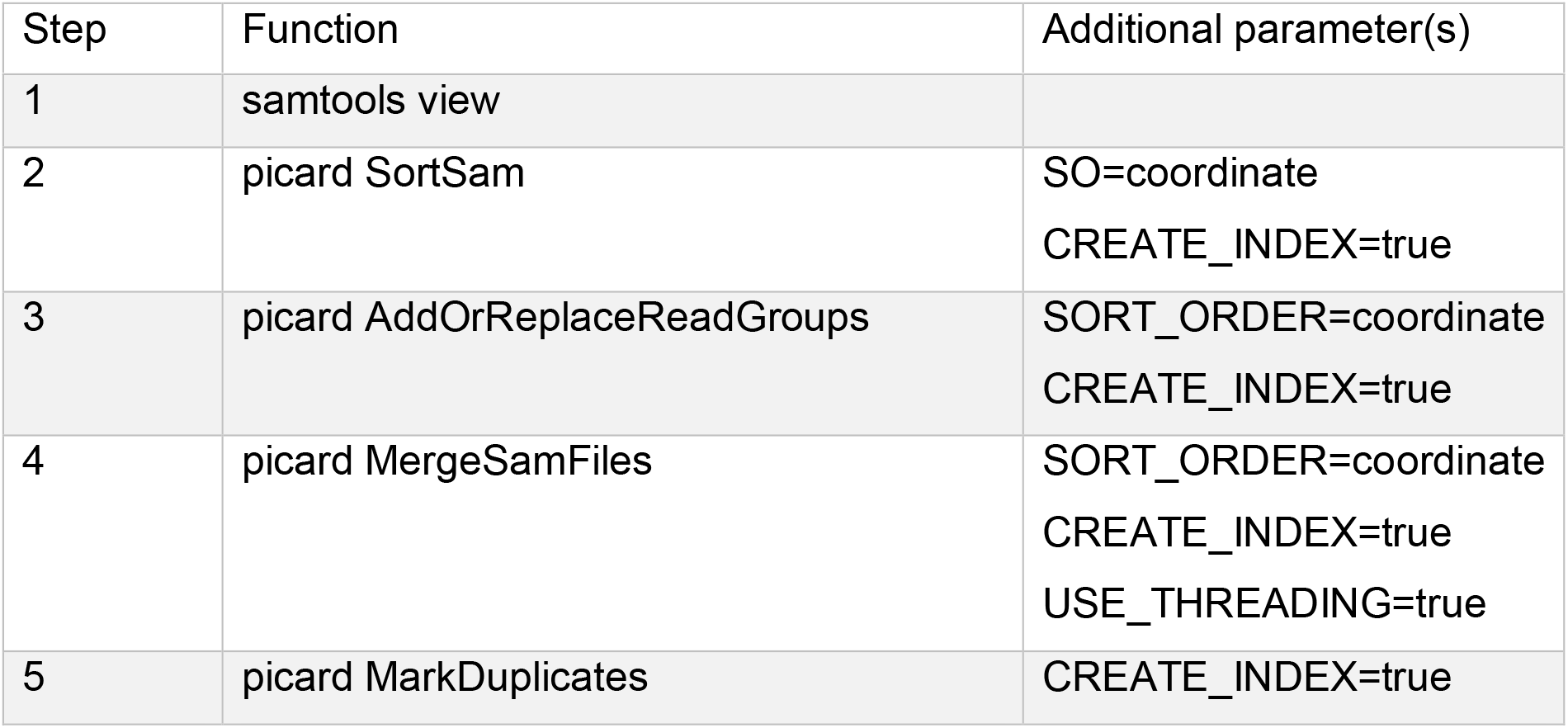
Explanation preprocessing steps. The various steps and functions used in this paper are explained *in this table, where the additional parameters are defined*.

| Step | Function | Additional parameter(s) |
| --- | --- | --- |
| 1 | samtools view |  |
| 2 | picard SortSam | SO=coordinate<br>CREATE_INDEX=true |
| 3 | picard AddOrReplaceReadGroups | SORT_ORDER=coordinate<br>CREATE_INDEX=true |
| 4 | picard MergeSamFiles | SORT_ORDER=coordinate<br>CREATE_INDEX=true<br>USE_THREADING=true |
| 5 | picard MarkDuplicates | CREATE_INDEX=true |

After the aforementioned steps listed in table 1, HTSeq^10^ is used to quantify the reads into count files (with the function htseq-counts) which are used in R algorithms for further analysis. Currently, single cell studies in BRAIN-SAT are processed from the raw count files that are provided by GEO.

### Bulk RNA analysis

After the preprocessing step, raw reads are filtered with the data-adaptive flag method for RNA-Seq data (DAFS)^12^. This method uses, a combination of the Kolmogorov-Smirnov statistics and multivariate adaptive regression splines to determine an optimal threshold value per sample to separate high and low expressed genes. The outcome of this filtering was saved in the MOLGENIS database.

The differential expression analysis function coverts the raw data with edgeR^12^, this analysis uses two selected conditions. The conditions are ordered alphabetically where condition “A” is used as a baseline condition (or control) and is compared to “B”, which is used as the condition of interest. This means that genes with a negative log fold change (logFC) are increased in the control condition. Whereas a positive logFC indicates an increase in the condition of interest. Differentially expressed genes are represented in an interactive scatterplot (generated with Plotly^13^) and is accompanied by a data table that consists of the gene symbol, logFC and false discovery rate (FDR). The data table can be used to find the logFC and FDR of the gene of interest, and the interactive scatterplot displays the overall differences between two conditions. These differences can be examined in more detail with the use of the zoom and hovering function that is available in the scatterplot.

The quantitative expression analysis uses transcript per million (TPM)^14^ to transform the data for the bar graph visualization (D3js^15^). TPM values show the number of transcription copies of a gene in a condition of interest in the selected studies.

### Single cell RNA analysis

Downloaded single cell transcriptome data sets were subjected to two filtering steps: first, cells with less than 500 expressed genes were identified as empty cells and are removed from the dataset. Second, the quantiles were calculated on expressed genes, where genes in the 25-quantile range were returned for further analysis. This matrix was used to obtain the top 100 most abundantly expressed genes per condition, filtering out low expressed genes. The number of cells per condition was adjusted if there were more than five different conditions available in the study. The top 10,000 genes were used for the downstream analysis (generating a matrix that consists of 500 columns and 10,000 rows).

A *SingleCellExperiment*^6^ object was created from the count per million (CPM) values, this object will be used as input for the rest of the process. The tSNE^5^ is calculated with the highest perplexity possible; this value is dependent on the dataset. Values of the tSNE are passed to a Vue component^17^ that calls on Plotly to generate an interactive plot. When a gene is searched, counts of the gene are obtained and returned to the Vue component which calculates the opacity for each individual dot (which represents a cell).

## 3 Results

### Basic features

BRAIN-SAT is a platform that was generated to facilitate interactive analyses of published RNA-Seq data from brain cells. This aim was accomplished by generating various visualizations based on analysis of the input data.

The homepage of BRAIN-SAT (figure 2) contains the following elements: (5) the search engine that visualizes the gene of interest in two datasets (one human and one mouse) and (6) the publications tab, where the datasets of processed studies can be found. This tab allows the performance of the DEA or QEA after selecting the study of interest in the available studies.

**Figure 2.**
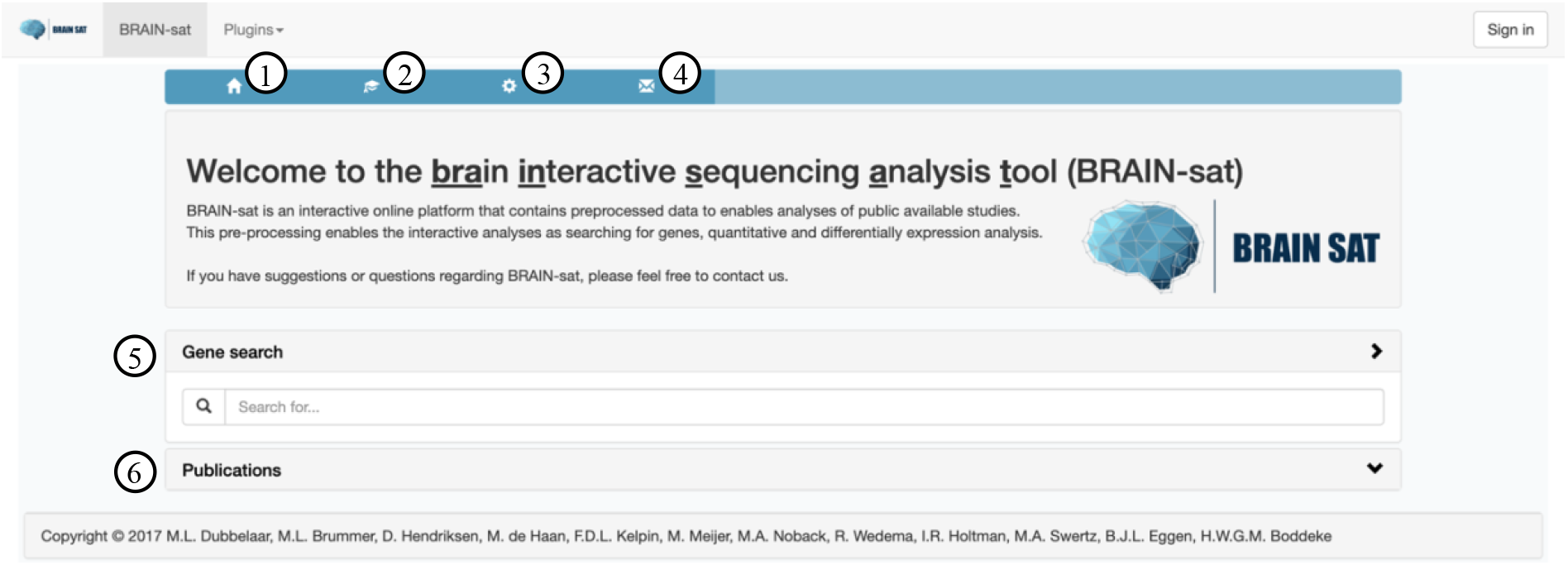
BRAIN-SAT homepage. Other pages can be accessed by using the available buttons: (1) homepage, (2) tutorial, (3) material and methods and (4) contact pages. The homepage of BRAIN-SAT contains two major functions: (5) the gene search bar and (6) the publications tab, which shows the available studies in BRAIN-SAT.

The homepage contains three buttons (see top left of the blue bar). The “home” button (1) is used to return to the home screen. An important part of the application is the tutorial page, which can be accessed by pressing the “education hat” icon (2). The “gear” button (3) redirects the user to the materials and methods page, which briefly explains the application that were used to create BRAIN-SAT and the workflow of the different analyses. The last button (the envelope) (4) leads to the contact information of the individuals that were most involved in the generation of BRAIN-SAT.

### Gene search

The BRAIN-SAT search engine on the homepage is used to depict the level of gene expression (log2(CPM)) based on different studies (cross-studies). An example of a search is represented in figure 3, where the gene *AXL* was used. The dot plot visualizes the median (or mean when only 2 replicates were available) expression of the control conditions per study. Whereas each different color is used to indicate a unique study and where each shape (dot, square or diamond) represents a different organism.

The dot plot in figure 3 shows the gene expression levels in several cell types, which are indicated on the x-axis. Hovering over a data point plot activates a text box that contains more information; the first line consists of the actual log2(CPM) value, the second line consists of the name of the first author and the year. Finally, the third line contains the region and if applicable the strain (which is identified between brackets).

**Figure 3.**
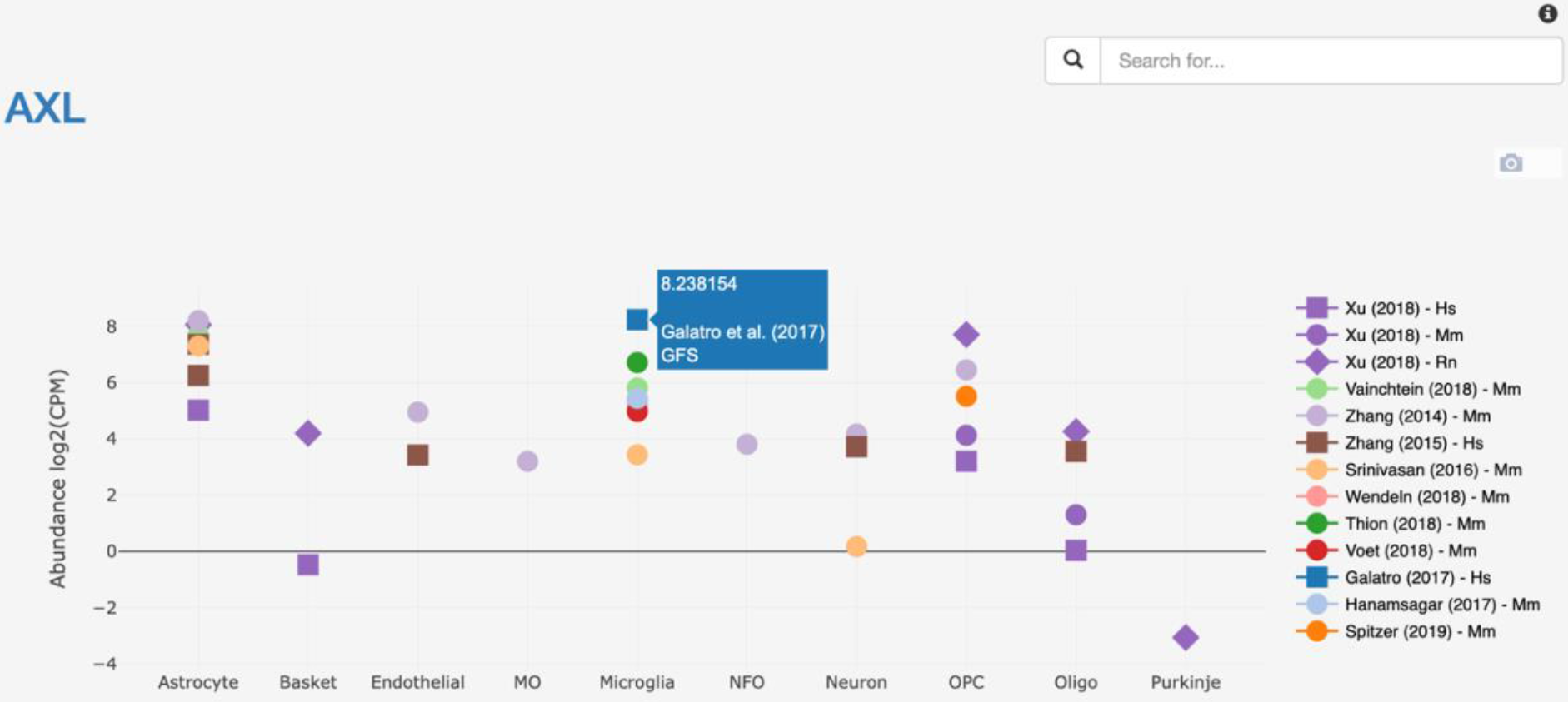
Cross-laboratory search of AXL. The x-axis depicts different cell types, both glia and neuronal subtypes. The y-axis shows the abundance (log2(CPM)) of AXL, which is indicated by the median (or mean) value that that is observed in the conditional samples. The color of the dot plots represents different studies (indicated by the author and year) and the shape of the data points (dot, square or diamond) represent the different organism.

### Quantitative expression analysis

The quantitative expression analysis (QEA) functionality can be accessed through the publication section, which consists of a collection of different studies. This analysis is done for each study separately, were different percentile ranges are used to describe the expression. The percentile description is ranked from not expressed to very high expressed (percentile range 0-5). For demonstration purposes the gene *Aldh1l1* (a gene that is used as a marker for astrocytes) was searched for in the dataset of Zhang *et al*. (2014). In figure 4 a moderately high expression (percentile range 10-20) is observed for the *Aldh1l1* gene in astrocytes. The error bar indicates the highest and lowest TPM values in relation to the median (or mean) of all samples, indicating that for *Aldh1l1*, the expression level of this gene was very consistent between the different samples.

**Figure 4.**
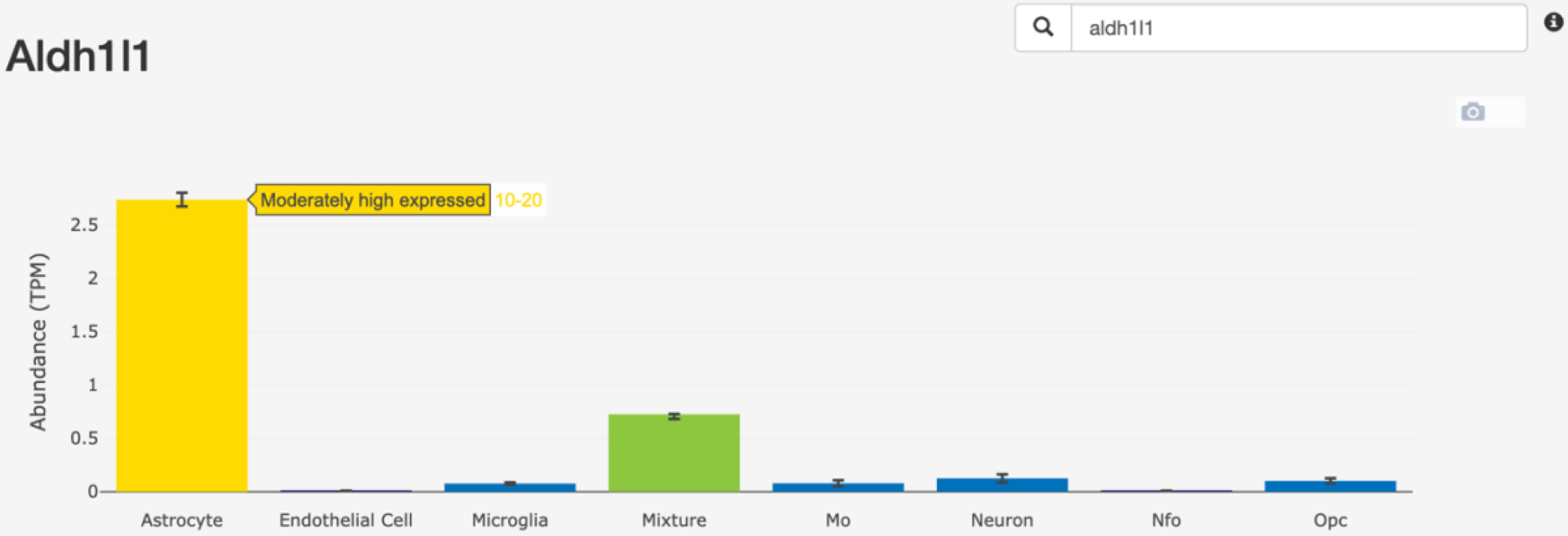
Aldh1l1 search (Zhang et al. 2014). The x-axis represents different conditions that are present in the study. The y-axis shows the gene expression (TPM). The colors of the bar plots can be used to indicate the expression level of the gene in a condition. A description of the percentile is shown when hovering over the bar plot.

### Differential expression analysis

The differential expression analysis (DEA) functionality enables an interactive pairwise comparison between two conditions to reveal changes in gene expression. Figure 5 shows the DEA between newly formed oligodendrocytes (NFO) and myelinating oligodendrocytes (MO). The volcano plot is divided into two parts (one for each condition), since the MO is used as baseline it can be found on the left side of the plot (logFC < −1, more highly expressed in MO). The right side of the plot shows genes that are more abundantly expressed in NFO cells (logFC > 1). The data table next to the plot depicts the −log10(FDR) and logFC of a gene of interest. A gene is only available in the table if the gene is significantly differentially expressed (FDR < 0.05 and an absolute logFC > 1).

**Figure 5.**
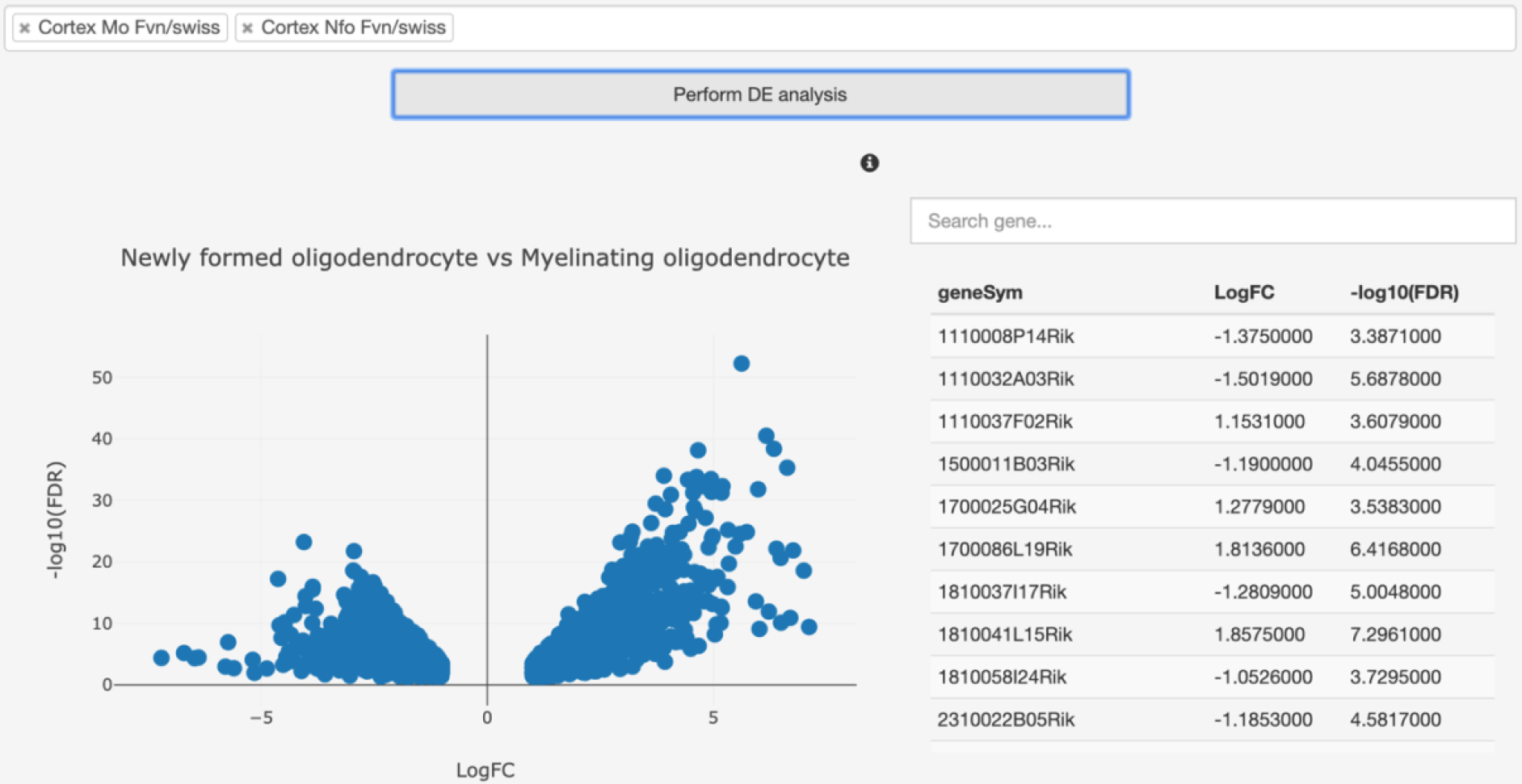
DEA NFO vs MO (Zhang et al. 2014). The left side of the page represents a volcano plot where the x-axis represents the logFC values and the y-axis the −log10(FDR) values. Each dot represents a differentially expressed gene, where the top-most right (NFO) or top left (MO) corners represent the most changed and significant differences between the conditions. The right side of the page contains a data table with genes that were found to be differentially expressed with their logFC and - log10(FDR) values.

### Single cell analysis

The newest feature is the analysis of single cell data which enables the visualization of the gene expression abundance in the cells with the highest expression level. The study of Matcovitch-Natan *et al*. is used in the interactive tSNE (figure 6A). Each dot represents a cell, and the color of the dot indicates a different condition. The x- and y-axis represent the two dimensions. The gene expression of *Irf8* can be observed in figure 6B with the use of three different visualizations to indicate the gene expression. The first visualization is an adaptation to the tSNE. The transparency of the dot is based on the abundance of gene expression in the cell. A more “solid” dot indicates high gene expression, whereas a transparent dot indicates low/no gene expression. Specific differences in gene expression based on the condition can be seen in the boxplot and pie card. The boxplot shows that the highest expression of *Irf8* can be observed in the condition “brain microglia E12.5”. The pie chart indicates that half of the detected *Irf8* expression derived from the “brain microglia E12.5” sample.

**Figure 6.**
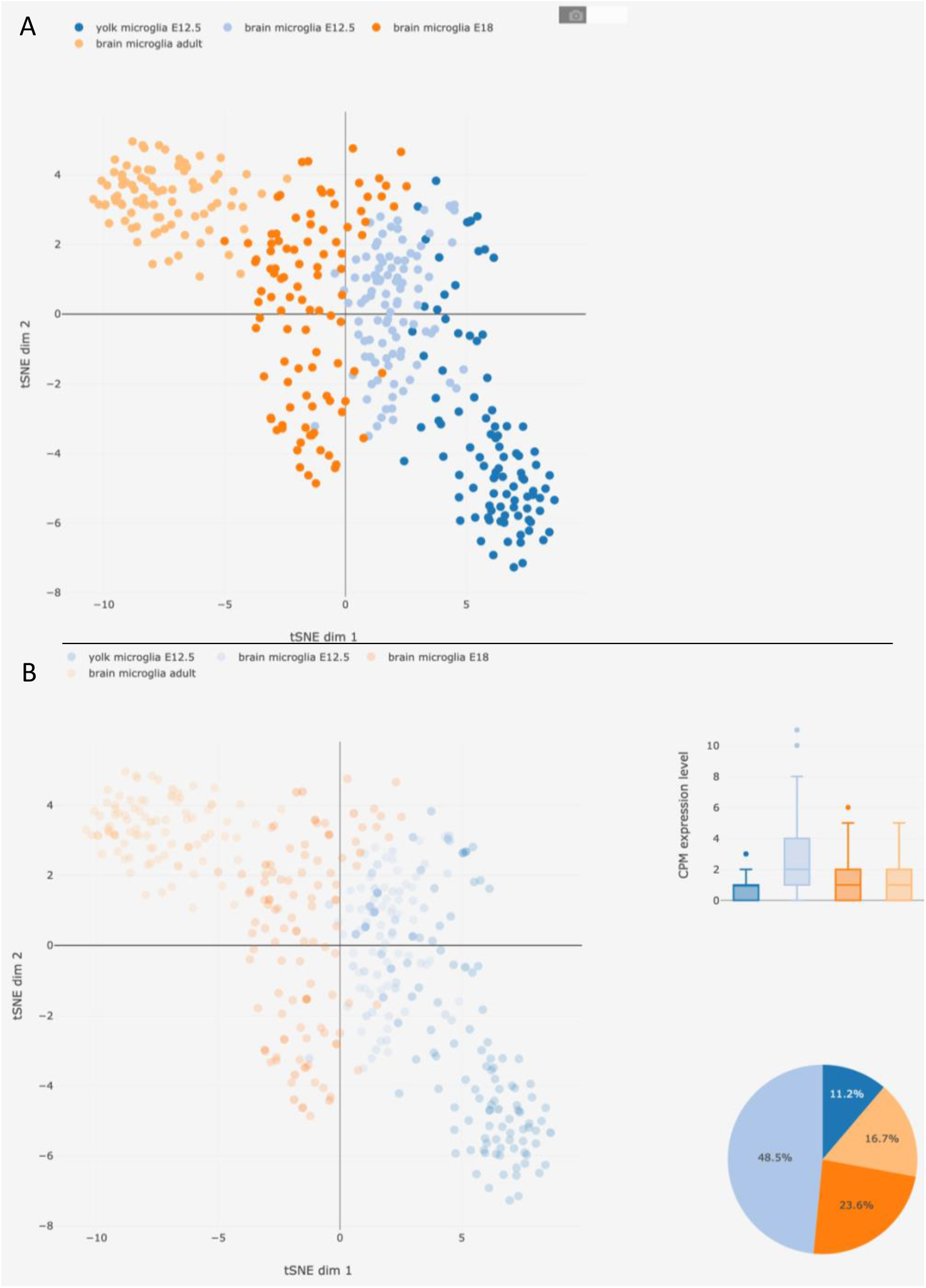
Single cell feature (Matcovitch-Natan et al. 2016). The output of the single cell analysis feature, which is divided into two parts. A) The tSNE is generated first, where the x- and y-axes represent the first and second dimension. Each dot represents a cell and the colors are used to distinguish the conditions. B) The dot transparency in the tSNE are altered after searching “Irf8” in the data of Matcovitch-Natan et al. (2016). The boxplot (top right) and the pie chart (bottom right) are used to indicate the expression per condition for “Irf8” in more detail.

### Comparison of BRAIN-SAT with other web applications

To date there are several applications available online that can be used to perform quantitative expression analysis. The most recent applications are: GOAD^3^, the Brain RNA-Seq application from Barres’s lab^18^, Neuroexpresso^19^ and the microglia single cell atlas^20^.These applications will be discussed below.

The aim of GOAD was to generate an accessible platform for glia biologists without the requirement of bioinformatic expertise. Information from various studies was aligned, quantified, and saved into a database. This information was used for visualization purposes. Studies available on the website cover different glia subtypes in different neurodegenerative diseases that could be used further for visualization through the expression analyses.

Brain RNA-Seq^18^ generated by Barres’s lab enables open access to the lab’s mouse and human data. The interface of the web application is easy to use. In addition, it is possible to access the quantitative expression several datasets that are publicly available on the website.

Neuroexpresso^19^ is described as a cross-laboratory database that combines data generated using GPL339 and GPL1261 micro-array chips together with a single cell RNA-Seq dataset that was generated by Tasic *et al*.^21^. In addition, only samples after postnatal day 14, wild-type and untreated animals were used. Neuroexpresso is able to identify gene markers that can be found in the homeostatic state of the various cell types.

The microglia single cell atlas by the lab of Stevens, contains single cell RNA-Seq data of microglia samples isolated at several ages across the lifespan from both female and male mice. In addition, the data includes microglia from saline and lysolecithin injected white brain matter. A visualization aspect is available through the search engine of the website.

## 4 Conclusion

The generation of BRAIN-SAT enabled the open accessibility of the data available from published studies. The FAIR principle was introduced to improve the available infrastructures and to enable data reusage^1^. We aimed to create an application that followed these principles when feasible. BRAIN-SAT consists of (meta)data that are globally unique, the data is open accessible and contains author and study information.

In addition, data is shared and could be further used for knowledge representation. Implementation of other principles might improve our application in the future, where more information about the samples and more high-quality data can be included in the near future.

Since the release of GOAD, several other applications became available for public use (table 4). These applications offer a range of different functionalities as: single cell and/or bulk RNA-Seq or cross-data analysis to explore available data. BRAIN-SAT was created to facilitate interactive transcriptome analyses, adding additional features such as performing a DEA on all available samples and exploring single cell RNA-Seq expression data. To gather datasets in the neuroscience field, we introduce a platform that researchers can use on open access data. The layout of BRAIN-SAT focusses on gene searches and enables open access to the data sets in the application, allowing faster turnover of the uploaded studies. This means that we can update the application faster, generating a more up to date application. The future perspectives for this application are as follows.

**Table 4:** Functionalities applications.

|  | GOAD | Brain RNA-Seq | NeuroExpresso | Microglia single cell | BRAIN-SAT |
| --- | --- | --- | --- | --- | --- |
| Gene search | x | x | x | x | x |
| QEA | x | x |  |  | x |
| DEA | x |  | x |  | x |
| Single cell analysis |  |  |  | x | x |
| Multiple organism | x | x |  |  | x |
| Cross library |  |  | x |  | x |
| Processing pipeline |  | ? | x |  | x |
| Interactive figures |  | x | x |  | x |
| Interactive analyses |  |  |  |  | x |
| Fast platform |  | x | x | x | x |

Fast incorporation of new studies in BRAIN-SAT, since data sets only need alignment and quantification. In addition, limiting the number of studies creates a collection of the most interesting findings in the field.

Adaptations of the website will be made based on new developments in the field. An example is the inclusion of the single cell analysis.

With the implementations of the above-mentioned perspectives we aim to generate an interactive platform that enables researchers to access published data and analyze it without extensive bioinformatic expertise.

## Acknowledgements

The creation of BRAIN-SAT would not have been possible without the Genomics Coordination Center in the UMCG. The knowledge regarding MOLGENIS, the transcriptomic pipelines and issues regarding the application and server were obtained through D. Hendriksen, M. de Haan, F.D.L. Kelpin, T. de Boer, B. Charbon, M.K. Slofstra, G. van der Vries, E. Adriaanse and M.A. Swertz and many colleagues of this department.

We would like to thank M.A. Noback and R. Wedema of the Hanze University of Applied Sciences for supervising both M.L. Dubbelaar and M.L. Brummer. Several aspects of BRAIN-SAT exist or are improved because their suggestions and guidance.

The authors would like to express their sincere gratitude to Dr. I.R. Holtman. BRAIN-SAT would not be the same without the provided guidance and input during the development of the glia open access database (GOAD). I.R. Holtman initiated the collaboration with the Genomics Coordination Center in the UMCG and the Hanze University of Applied Sciences.

## References

1. Wilkinson MD, Dumontier M, Aalbersberg IjJ, et al. The FAIR Guiding Principles for scientific data management and stewardship. Sci Data. 2016;3:160018. doi:10.1038/sdata.2016.18

2. Manzoni C, Kia DA, Vandrovcova J, et al. Genome, transcriptome and proteome: the rise of omics data and their integration in biomedical sciences. Brief Bioinform. 2018;19(2):286–302. doi:10.1093/bib/bbw114

3. Holtman IR, Noback M, Bijlsma M, et al. Glia Open Access Database (GOAD): A comprehensive gene expression encyclopedia of glia cells in health and disease. Glia. 2015;63(9):1495–1506. doi:10.1002/glia.22810

4. Swertz MA, Dijkstra M, Adamusiak T, et al. The MOLGENIS toolkit: rapid prototyping of biosoftware at the push of a button. BMC Bioinformatics. 2010;11(Suppl 12):S12. doi:10.1186/1471-2105-11-S12-S12

5. van der Maarten LJP, Hinton G. Visualizing Data using t-SNE. J Mach Learn Res. 2008;9:2579–2605. http://www.jmlr.org/papers/volume9/vandermaaten08a/vandermaaten08a.pdf.

6. Edgar R, Domrachev M, Lash AE. Gene Expression Omnibus: NCBI gene expression and hybridization array data repository. Nucleic Acids Res. 2002;30(1):207–210. http://www.ncbi.nlm.nih.gov/pubmed/11752295.

7. EMBL-EBI. European Nucleotide Archive. https://www.ebi.ac.uk/ena. Published 2019.

8. Li H, Handsaker B, Wysoker A, et al. The Sequence Alignment/Map format and SAMtools. Bioinformatics. 2009;25(16):2078–2079. doi:10.1093/bioinformatics/btp352

9. Broadinstitute. Picard. 2016. http://broadinstitute.github.io/picard/.

10. Anders S, Pyl PT, Huber W. HTSeq--a Python framework to work with high-throughput sequencing data. Bioinformatics. 2015;31(2):166–169. doi:10.1093/bioinformatics/btu638

11. George NI, Chang C-W. DAFS: a data-adaptive flag method for RNA-sequencing data to differentiate genes with low and high expression. BMC Bioinformatics. 2014;15(1):92. doi:10.1186/1471-2105-15-92

12. Robinson MD, McCarthy DJ, Smyth GK. edgeR: a Bioconductor package for differential expression analysis of digital gene expression data. Bioinformatics. 2010;26(1):139–140. doi:10.1093/bioinformatics/btp616

13. Plotly Technologies Inc. Collaborative data science Publisher: Plotly Technologies Inc. 2015.

14. Li B, Ruotti V, Stewart RM, Thomson JA, Dewey CN. RNA-Seq gene expression estimation with read mapping uncertainty. Bioinformatics. 2010;26(4):493–500. doi:10.1093/bioinformatics/btp692

15. Bostock M, Ogievetsky V, Heer J. D3: Data-Driven Documents. 2011. https://github.com/d3/d3.

16. Lun A, Risso D. SingleCellExperiment: S4 Classes for Single Cell Data. 2019.

17. You E. Vuejs. 2013.

18. Barres Lab. Brain RNA-Seq.

19. Mancarci BO, Toker L, Tripathy SJ, et al. Cross-Laboratory Analysis of Brain Cell Type Transcriptomes with Applications to Interpretation of Bulk Tissue Data. eneuro. 2017;4(6):ENEURO.0212-17.2017. doi:10.1523/ENEURO.0212-17.2017

20. Hammond TR, Dufort C, Dissing-Olesen L, et al. Single-Cell RNA Sequencing of Microglia throughout the Mouse Lifespan and in the Injured Brain Reveals Complex Cell-State Changes. Immunity. 2018.

21. Tasic B, Menon V, Nguyen TN, et al. Adult mouse cortical cell taxonomy revealed by single cell transcriptomics. Nat Neurosci. 2016;19(2):335–346. doi:10.1038/nn.4216

